# Statistical Pattern Recognition Reveals Shared Neural Signatures for Displaying and Recognizing Specific Facial Expressions

**DOI:** 10.1101/2019.12.15.873737

**Authors:** Sofia Volynets, Dmitry Smirnov, Heini Saarimäki, Lauri Nummenmaa

## Abstract

Human neuroimaging and behavioural studies suggest that somatomotor “mirroring” of seen facial expressions may support their recognition. Here we show that viewing specific facial expressions triggers the representation corresponding to that expression in the observer’s brain. Twelve healthy female volunteers underwent two separate fMRI sessions: one where they observed and another where they displayed three types of basic facial expressions (joy, anger and disgust). Pattern classifier based on Bayesian logistic regression was trained to classify facial expressions i) within modality (trained and tested with data recorded while observing or displaying expressions) and ii) between modalities (trained with data recorded while displaying expressions and tested with data recorded while observing the expressions). Cross-modal classification was performed in two ways: with and without functional realignment of the data across observing / displaying conditions. All expressions could be accurately classified within and also across modalities. Brain regions contributing most to cross-modal classification accuracy included primary motor and somatosensory cortices. Functional realignment led to only minor increases in cross-modal classification accuracy for most of the examined ROIs. Substantial improvement was observed in the occipito-ventral components of the core system for facial expression recognition. Altogether these results support the embodied emotion recognition model and show that expression-specific somatomotor neural signatures could support facial expression recognition.

## Introduction

Humans convey their internal states, motives and needs with facial expressions. So-called basic emotion theories propose that the evolution has carved out discrete neural circuits and physiological systems that support distinct survival functions (Panksepp 1982; Ekman 1992), and that activation of each discrete system would be associated with specific pattern of facial muscle activity resulting in emotion-specific facial expressions (Ekman 1999; for a more recent formulation, see Keltner et al., 2019). These expressions are used for communication in social interaction, as they convey information about the expressers’ internal states that can be “read out” from the face. Expression recognition is however not carried only by the brain’s visual systems (Haxby et al. 2000). Models of embodied emotion recognition (see reviews in Niedenthal 2007 and Wood et al. 2016) have proposed that sensorimotor simulation of seen facial expressions could support their recognition, especially when the recognition cannot be achieved via the straightforward automated visual pattern-recognition strategies (Winkielman et al. 2015; Wood et al. 2016).

In line with the embodiment hypothesis, seeing others’ facial expressions triggers automatic facial mimicry (see e.g. Dimberg and Thunberg 1998). Neuroimaging studies have also established that displaying and observing facial expressions activates overlapping brain regions including premotor and somatosensory cortices (Carr et al. 2003; Hennenlotter et al. 2005; van der Gaag et al. 2007; Kircher et al. 2012). Furthermore, both damage to somatosensory cortex (Adolphs et al. 2000) and its inactivation by transcranial magnetic stimulation (TMS; Pourtois et al. 2004) impairs recognizing emotions from facial expressions, suggesting that somatomotor embodiment of seen emotions supports their recognition. In line with these results, emotional facial expressions can be successfully decoded from motor brain regions (Liang et al., 2017). Yet strong support for the embodied recognition view would require showing that i) both displaying and seeing different facial expressions would trigger *expression-specific*, discrete neural signatures in the somatomotor system and that ii) these expression-specific neural signatures would be *corresponding* when displaying and observing the expressions. Previous studies have found that different seen facial expressions elicit discernible neural activation patterns in visual areas and multisensory temporal cortical areas (e.g. Said et al. 2010; Harry et al. 2013; Peelen et al., 2010; Wegrzyn et al. 2015), but it remains unresolved whether these and somatomotor activation patterns converge with those elicited while displaying specific facial expressions.

Here we tested the embodied emotion recognition model directly by using functional magnetic resonance imaging (fMRI) and statistical pattern recognition techniques. Participants observed and displayed three types of basic facial expressions (joy, anger and disgust) while their brain activity was measured with fMRI. Subsequently, a pattern classifier based on Bayesian logistic regression was trained to classify the observed or displayed emotions from the fMRI signals using regions-of-interest (ROIs) in the face perception system (Haxby et al. 2000), emotion circuit (Saarimäki et al. 2016), and visual and somatomotor areas. The critical test involved training the classifier with data recorded while displaying expressions and testing with data recorded while observing the expressions. Successful classifier performance in this condition, and particularly when using data from the somatosensory and motor cortices, would provide support for expression-specific embodiment during perception of facial expressions. We show that different facial expressions have distinguishable somatomotor neural signatures that are activated similarly both when viewing and displaying the expressions.

## Materials and Methods

### Participants

Twelve healthy right-handed female volunteers (mean age 21 years, range 20–26 years) with normal or corrected to normal vision and normal hearing (self-reported) volunteered for the study. None had a history of neurological or psychiatric diseases, or current medication affecting the central nervous system. All subjects signed informed consent forms, approved by the Aalto University Institutional Review Board, and were compensated for their time. All subjects were pre-tested for their ability to recognize emotional facial expressions. In this recognition test, subjects viewed photographs of unfamiliar models displaying facial expressions of basic emotions (anger, fear, disgust, happiness, sadness, and surprise) as well as morphed versions (30% and 60% morphs with neutral expressions) of the same photos. The test stimuli were derived from the Karolinska Directed Emotional Faces database (KDEF; Lundqvist et al. 1998). Subjects were asked to identify emotion on the given photo in a 6-alternative forced choice test. Mean accuracy was 74% and exceeded chance level (16.6%) for all expressions.

### Experimental Design and Statistical Analysis

#### Training phase

At least one day prior to the fMRI experiment all subjects participated in an individual training session, where the experimental setup and tasks were explained. Main facial features of the expressions of joy, anger, and disgust were described according to the Facial Action Coding System (FACS; Ekman and Friesen 1978). Participants were then instructed to 1) select one triggering memory/image for eliciting each emotion (joy, anger, disgust) to make the expressions more genuine (e.g. favourite joke to elicit smile) and 2) rehearse displaying the corresponding facial expressions when prompted with minimal head motion.

#### Displaying Expressions Task

Experimental design is summarized in Fig 1. During the Displaying Expressions Task (Fig 1a) participants displayed facial expressions of joy, anger and disgust while being scanned with fMRI. Each trial started with an auditory instruction, specifying the facial expression to be displayed (spoken words “joy”, “anger”, or “disgust”). Next, a beep indicated that the subject should display the facial expression and keep the full-blown pose until a second beep (after 5 s) indicated the end of the trial. During the 10-15 s intertrial interval the subject was instructed to relax their face. A fixation cross was shown at the centre of the screen throughout the experiment. The Displaying part consisted of 4 runs with 24 trials in each, and the subjects displayed each expression 8 times per each run.

**Figure 1.**
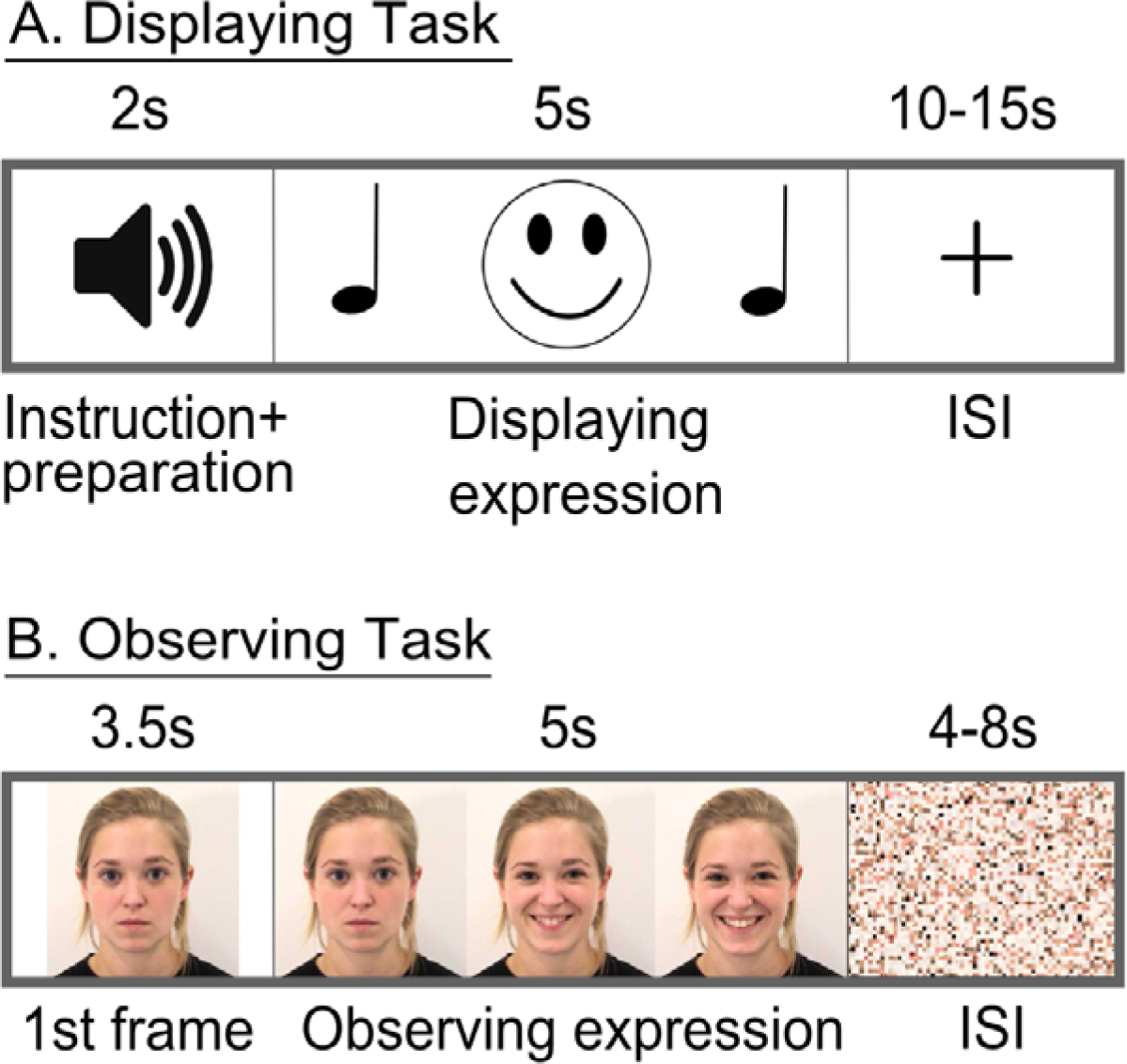
Experimental design. (A) In the Displaying task, participants heard a word describing the facial expression to be performed next. After 2 seconds they heard a beep sound, which marked the start of expression execution. During the next 5 seconds, participants went from neutral face to the fully blown target facial expression and kept it until the second auditory cue which prompted the participants to relax their facial muscles for the subsequent 10-15 s interstimulus interval (ISI). (B) In the Observing task, participants were first shown a neutral facial expression of the model from upcoming video for 3.5 s. After that, they viewed a 5s dynamic facial expression video, where the model went from neutral face to fully blown emotional expression and kept it until the end of the video. During the 4-8 s ISI, a scrambled facial expression was shown.

#### Observing Expressions Task

In the Observing Expressions Task (Fig 1b) participants viewed short video clips (5 s) of six models (three females) displaying facial expressions of joy, anger and disgust. The stimuli were selected from ADFES database (Van der Schalk et al. 2011), and the models in the clips were unfamiliar to the participants. All clips begun with a neutral face, followed by a dynamic display of the facial expression. Prior to each clip, subjects were shown the 1^st^ frame of the video (i.e. neutral face) for 3.5s. This was followed by the dynamic expression, that was held in its full-blown phase until the end of the clip. Each stimulus was followed by a random 4-8s rest period. To avoid peaks in low-level visual cortical activations, a scrambled picture of the upcoming model was shown during the rest period.

To keep participants focused on the task, three trials per run contained a still picture of the neutral face instead of the video clip. Participants were asked to press the response button as soon as they detected a trial without any facial motion. These trials were excluded from the analysis. The Observing part consisted of 4 runs with 24+3 trials in each, and the subjects observed the videos of each facial expression 8 times per each run.

#### Stimulus presentation

Stimulus delivery was controlled using Presentation software (Neurobehavioral Systems Inc., Albany, CA, USA). Visual stimuli were back-projected on a semi-transparent screen using a 3-micromirror data projector (Christie X3, Christie Digital Systems Ltd., Mönchengladbach, Germany) and reflected via a mirror to the subject. Auditory cues were delivered with Sensimetrics S14 insert earphones (Sensimetrics Corporation, Malden, MA, United States). Sound intensity was adjusted for each subject to be loud enough to be heard over the scanner noise. Three subjects were also recorded with an fMRI-compatible face camera to ensure that the facial expressions were displayed successfully during the experiment.

#### fMRI acquisition and preprocessing

MRI scanning was performed on two sessions on separate days. The data were acquired with 3T Siemens Magnetom Skyra scanner at the Advanced Magnetic Imaging Centre, Aalto NeuroImaging, Aalto University, using a 20-channel Siemens head coil. Whole-brain functional images were collected using a whole brain T2*-weighted echo-planar imaging (EPI) sequence, sensitive to blood oxygenation level-dependent (BOLD) signal contrast, with the following parameters: 33 axial interleaved slices, TR = 1.7 s, TE = 24 ms, flip angle = 70°, voxel size = 3 x 3 x 4.0 mm3, matrix size = 64 x 64 x 33. A total of 245 volumes were acquired in each run for the Observing task, and 275 volumes were acquired in each run for the Making task. The first 5 volumes of each run were discarded. High-resolution anatomical images with isotropic 1 mm^3^ voxel size were collected using a T1-weighted MP-RAGE sequence.

The fMRI data were preprocessed using MATLAB (The MathWorks, Inc., Natick, Massachusetts, USA), FSL (FMRIB’s Software Library, www.fmrib.ox.ac.uk/fsl) and SPM8 (www.fil.ion.ucl.ac.uk/spm/). After slice timing correction, the functional images were realigned to the middle scan by rigid-body transformations with MCFLIRT to correct for subject motion. Non-brain matter was next removed using BET (Smith 2002). Functional images were then registered to the MNI152 standard-space template (Montreal Neurological Institute) with 2-mm resolution. The transformation parameters were acquired by first calculating transformations from structural to standard space and from functional to structural space, and then concatenating these parameters. Next, these transformation parameters were used to co-register functional datasets to the standard space. Both registration steps were performed using FLIRT (Jenkinson et al. 2002). Motion artefacts were cleaned from the functional data using 24 motion regressors (Power et al. 2014). None of the subjects were excluded from analysis due to excessive head motion during the Displaying task, as framewise displacement values (Power et al. 2012) exceeded 0.5mm in less than 1.81% of all timepoints per subject. Signal from white matter, ventricles and cerebrospinal fluid were cleaned from the data as implemented in BraMiLa pipeline (https://git.becs.aalto.fi/bml/bramila). For the general linear model (GLM) analysis, additional spatial smoothing step with a Gaussian kernel of FWHM 8 mm was also applied; the subsequent classification analysis was run on unsmoothed data.

#### Task-evoked BOLD responses

Task-evoked responses to execution and observation of facial expressions were analysed using the two-stage random effects analysis with GLM implemented in SPM8 (www.fil.ion.ucl.ac.uk/spm). Boxcar regressors (displaying and observing facial expressions) were used to model fMRI voxel time series. The regressors included the time points when the facial expressions were observed or displayed, respectively. Regressors were convolved with the canonical hemodynamic response function (HRF) to account for hemodynamic lag. The input data were high-pass filtered with 128-s cut-off. After generating subject-wise contrast images, a second level (random effects) analysis was applied to these contrast images in a new GLM to allow population-level inference. Statistical threshold was set at *p* < .05, false discovery rate (FDR) corrected at cluster level.

#### Region-of-interest selection

For the classification analysis, four sets of ROIs were employed. First, we used the whole grey matter (derived from MNI standard brain template; Grabner et al. 2006) as a single ROI, and functional ROIs based on the experimental tasks. The ROIs were derived from the group level activations for a) observing and b) displaying facial expressions, c) their intersection and d) their union. The contrast images from second-level GLM analysis were thresholded at T > 2 for Displaying task and at T > 3 for Observing task. Liberal threshold was chosen as the maps were not used for statistical inference, but rather as a priori feature-selection filter that would capture the expression display- and observation-dependent neural activation. Second, we used anatomically defined ROIs in the emotion, somatosensory, and face perception circuits defined using the Harvard-Oxford cortical and subcortical structural atlases (Desikan et al. 2006). The emotion circuit ROIs included insula, thalamus and anterior cingulate cortex (ACC; Saarimäki et al. 2016). The somatomotor ROIs included primary somatosensory cortex (SI), secondary somatosensory cortex (SII), primary motor cortex, and premotor cortex. The face perception circuit ROIs included the key nodes of the core system for face perception (Haxby et al. 2000): inferior occipital cortex, fusiform cortex, superior temporal sulcus (STS), and MT/V5 region.

#### Pattern classification

Pattern classification was performed in three ways. First, we wanted to test whether each of the displayed and observed expressions is associated with distinct neural signatures. To that end, we performed a conventional within-modality classification, where the classifier was trained and tested with data from the same modality (displaying or observing). Second, to test whether displaying and observing facial expressions would be associated with similar, expression-specific neural signatures in the brain, we initially performed cross-modal classification without functional realignment. In this approach, the data from the Displaying condition were used to train the pattern classifier to distinguish between the three different expressions, and then the classifier was validated using corresponding data from the Observing condition. Third, to test whether the neural codes for observing and displaying facial expressions would be similar, yet anatomically misaligned, we performed cross-modal classification analysis where an additional realignment step was employed to allow functional coregistration of Observing and Displaying data.

For all tested classifiers, the data comprised all trials, with 3 TRs per trial, recorded during displaying and observing phases of the experiment, and shifted by 6 s to account for the hemodynamic lag, with each TR independently used as a training or testing example. For all tested classifiers, we evaluated the performance of the classification model in leave-one-run-out cross-validation framework, where three runs were used to train the classifier and the left-out run was used in testing, and the process was repeated iteratively for each run. In cross-modal classification analyses, the training runs were taken from Displaying data, and the testing run was taken from the Observing data because we specifically wanted to test whether of expression display-related activity would be predictive of the seen facial expressions.

Classification was performed with Bayesian logistic regression with a sparsity promoting Laplace prior to classify brain activity patterns measured during displaying and observing facial expressions (van Gerven et al. 2010). Each individual voxel weight was given a univariate Laplace prior distribution with a common scale hyperparameter to promote sparsity in the posterior distribution (Williams 1995). The multivariate posterior distribution was approximated using the expectation propagation algorithm (van Gerven et al. 2010) implemented in the FieldTrip toolbox (Oostenveld et al. 2011). Four binary classifiers were trained to discriminate between each expression category versus the others. The classification performance was tested by collecting the class probabilities for each pattern in the testing set using the binary classifiers and assigning the class with the maximum probability to each pattern.

#### Functional realignment and cross-modal classification

To test for possible differences in regional organization of displaying and observing facial expressions, an additional functional realignment step was introduced in the analysis pipeline (see Smirnov et al, 2017, for details and validation of the approach). Briefly, we used Bayesian canonical correlation analysis (BCCA; Klami et al. 2013) to perform the realignment step prior to cross-modal classification. BCCA was implemented using R CCAGFA package (Virtanen et al. 2012; Klami et al. 2013). The BCCA model separates the correlation patterns in the simultaneous brain-activity spaces of display and observation of the same facial expression into three types of components: display-specific, observation-specific, and shared. The shared components provide a low-rank linear mapping for the realignment of the brain-activity spaces. Classifier was trained on the displaying data from the three runs and tested on functionally realigned observation data run.

#### Significance testing for classifier performance

Statistical significance of mean classification accuracies was tested by comparing them against chance level. Since empirical chance level can differ from naïve chance level (Combrisson and Jerbi 2015), we estimated chance level by randomly shuffling the class labels in 100 permutations. The accuracies of within- and cross-modal classification, as well as cross-modal classification after realignment were approximately normally distributed; hence the confidence intervals for their means were obtained from Student’s t-distributions with variance pooled across individuals.

## Results

### Task-related BOLD responses

Figure 2 shows brain regions activated during Displaying and Observing tasks in the main experiment (all expressions combined). Both Displaying and Observing tasks engaged precentral gyrus (motor strip), supramarginal gyrus, supplementary motor area (SMA), superior and inferior frontal gyri, temporal pole, as well as parietal operculum (S2), caudate, insula, and right thalamus. In general, observing versus displaying facial expressions yielded more widespread activity, yet 50% of the voxels activated during the Displaying task were also activated during the Observing task. Significant observation-specific activations were found in frontal pole, putamen, amygdala, lateral occipital cortex, parahippocampal, superior temporal, and inferior temporal gyri. The only area uniquely activated during the Displaying task was ACC. Additionally, somatomotor responses (precentral and postcentral gyri, SMA) were more widespread in the Displaying versus Observing task.

**Figure 2.**
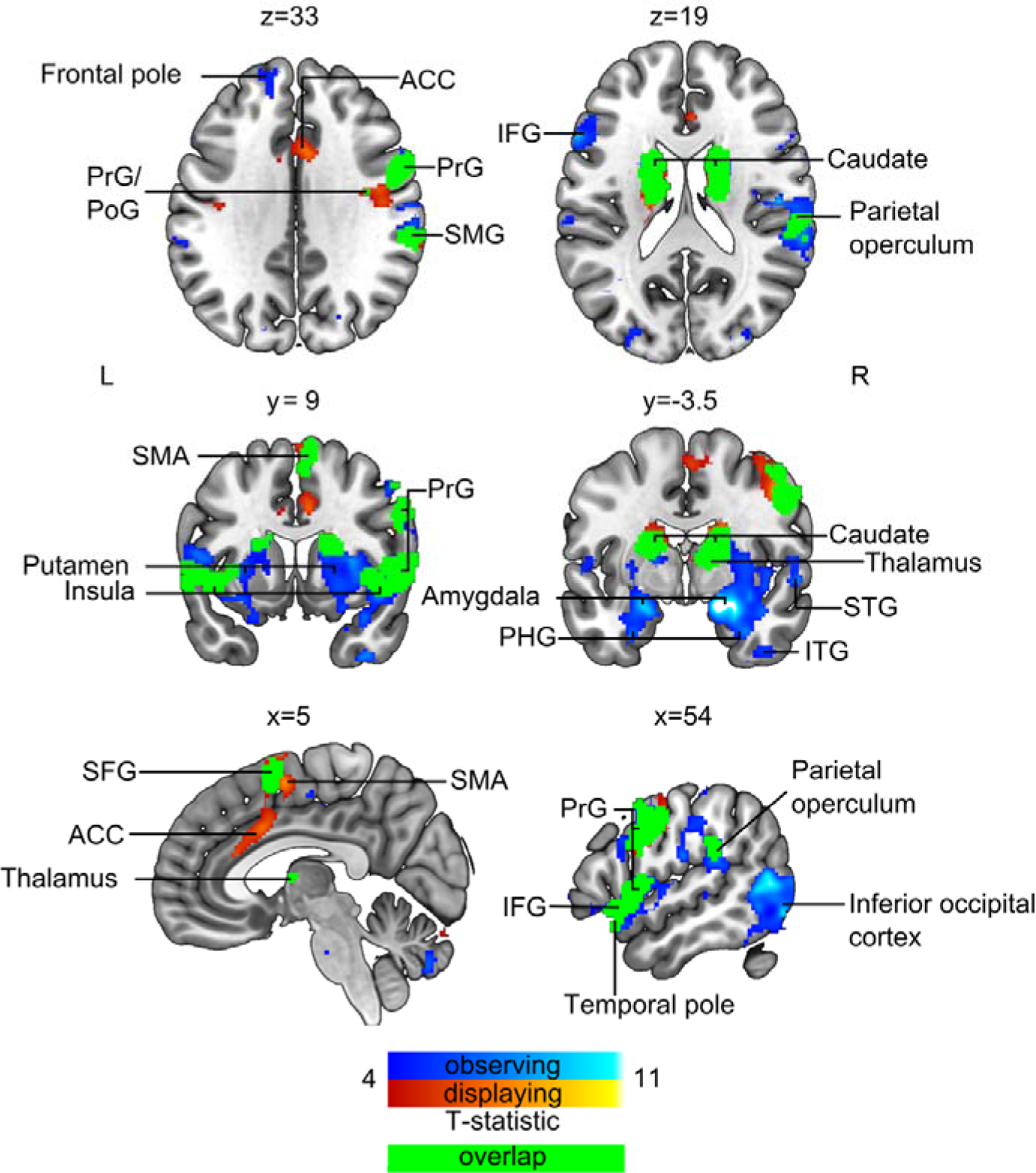
Brain regions responding to displaying (hot colours) and observing (cool colours) facial expressions. Green colouring shows the overlap between displaying and observing dependent activations. The data are thresholded at p < 0.05, FDR corrected. ACC – anterior cingulate cortex, PrG – precentral gyrus, PoG – postcentral gyrus, SMG – supramarginal gyrus, SMA – supplementary motor area, SFG – superior frontal gyrus, IFG – inferior frontal gyrus, STG – superior temporal gyrus, ITG – inferior temporal gyrus, PHG – parahippocampal gyrus.

### Within-modality classification for displaying and observing facial expressions

First, we ran conventional within-modality pattern classification analyses to test whether the three displayed or observed facial expressions have distinct neural signatures. For the Displaying condition, all functional and anatomical ROIs yielded statistically significantly above chance level accuracy (Fig 4–1 and **Fig S-1**). Mean accuracies varied from 39% to 64% against 33% chance level (*Ws* = 78, *ps* < .001). Accuracy was highest for whole grey matter and Displaying-Observing union ROIs (64%), followed by Displaying (62%) and Observing (61%) ROIs. Importantly, all facial expressions could be classified from each other significantly above chance level (Fig 3a). Within-modality classifier for Observing condition also yielded above chance level accuracy in all ROIs, except thalamus, with accuracies ranging from 36 to 54% against 33% chance level (*Ws* = 61-78, *ps* < .05; Fig 4–1). Accuracy was highest for inferior occipital cortex (54%), followed by whole grey matter (52%) and Displaying-Observing union (51%) ROIs. Again, all facial expressions could be classified from each other significantly above chance level (Fig 3b). Within-modality classification was significantly more accurate for displaying versus observing emotions in all ROIs except for inferior occipital cortex, V5 and ACC (*Ws* = 59-78, *ps* < .05).

**Figure 3.**
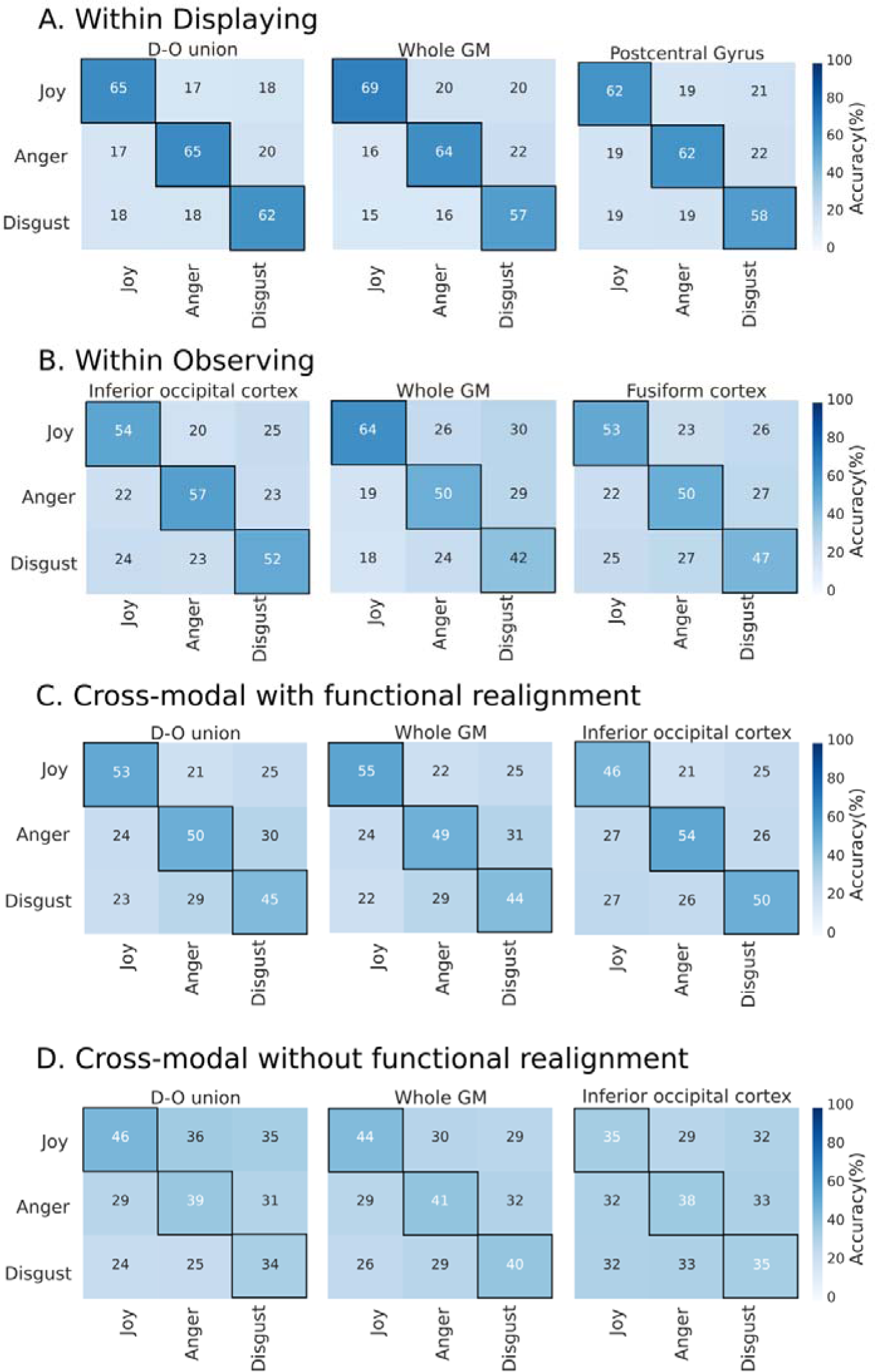
Confusion matrices for within-modality (A-B) and cross-modal classification with functional realignment (C) in representative regions of interest. All accuracies in the main diagonal significantly exceed chance level.

**Figure 4.**
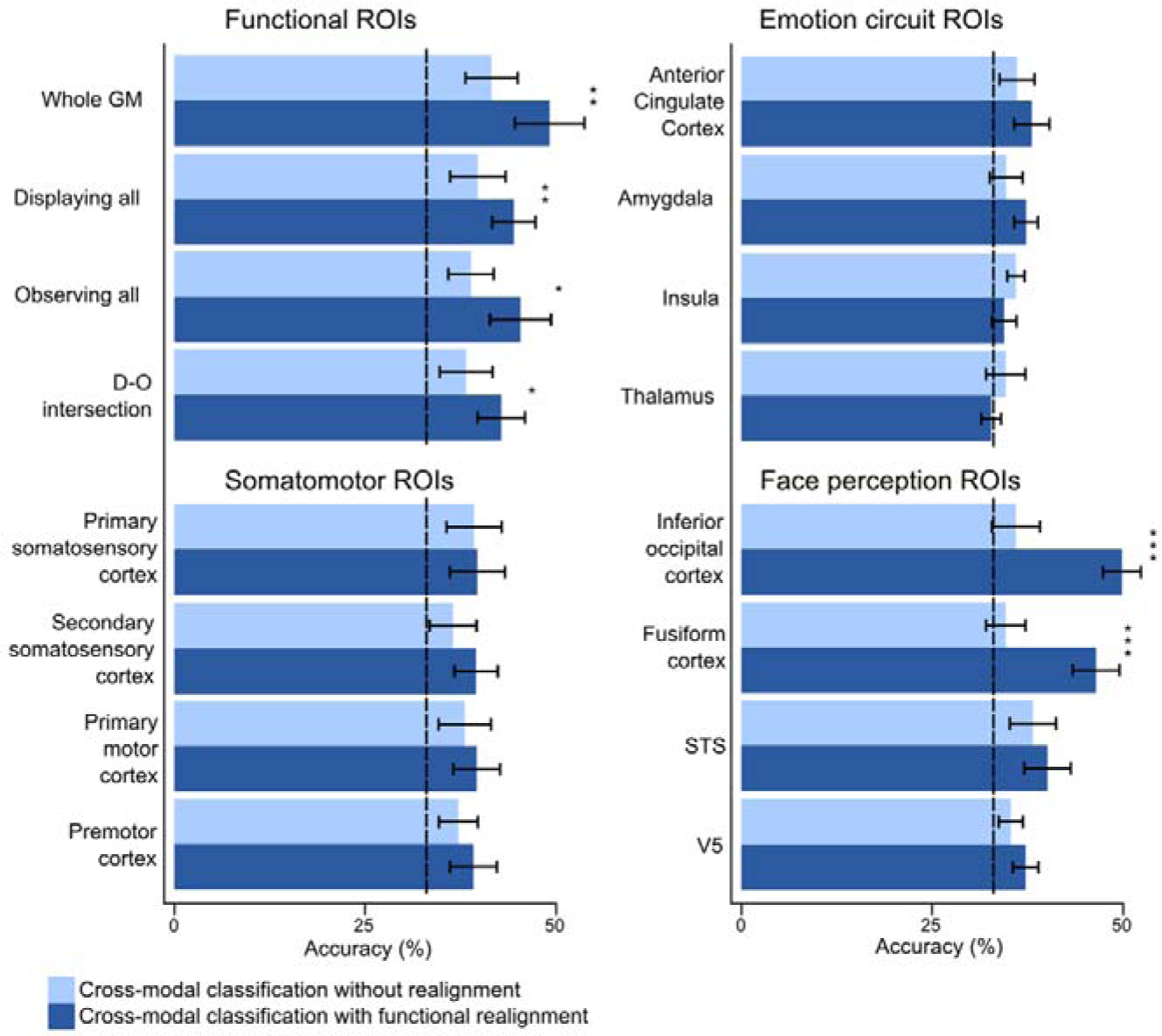
Mean classification accuracies (in %) and 95% confidence intervals for cross-modal classification with and without functional realignment in functional ROIs as well as in key components in the emotion, somatomotor, and face perception related regions. Dashed line represents the 33% chance level. Asterisks reflect statistical significance of the difference between classification accuracies before and after functional realignment of the data accordingly to the results of Wilcoxon tests (*** p < .001, ** p < .01, * p < .5). D-O intersection – intersection of Displaying expressions and Observing expressions ROIs derived from experimental data, STS – superior temporal gyrus, V5 – visual cortex area V5.

### Cross-modal classification with and without functional realignment

Accuracy of cross-modal classification exceeded the chance level already before functional realignment in most of the ROIs (*Ws* = 62-76, *ps* < .05; Fig 4 and Fig 4–1). Accuracy was highest in the whole grey matter ROI (42%), followed by Displaying-Observing union (40%) and Displaying ROI (40%). Chance level was not exceeded significantly for thalamus (35%), fusiform cortex (35%), amygdala (35%), and inferior occipital cortex (36%). After functional realignment, classification accuracy improved statistically significantly in the inferior occipital cortex, fusiform cortex, whole grey matter and functional ROIs derived from experimental data (*Ws* = 54-78, *ps* < .05, mean increase 2-14%; Fig 4). Accuracy was highest in inferior occipital cortex (50%), whole grey matter (49%), Displaying-Observing union (49%) and fusiform cortex (46%) ROIs, followed by the rest of functional ROIs with accuracies ranging from 43 to 45% (*Ws* = 77-78, *ps* < .001) and anatomically defined ROIs with accuracies ranging from 37 to 40% (*Ws* = 73-78, *ps* < .01). After functional realignment, chance level was not exceeded significantly only for thalamus (33%) and insula (34%).

Finally, we validated that neural responses to displaying and observing of each individual expression could be classified across modalities with approximately the same precision. After functional realignment, the classifier was able to distinguish expressions from each other with accuracies comparable across the ROIs: disgust 44-50%, anger 49-54%, and joy 46-55% (Fig 3c). The accuracy ranges for cross-modal classification before functional realignment were considerably lower: disgust 31-40%, anger 33-41%, and joy 29-46% (Fig 3d). However, the best discrimination was achieved in within-modality classification on Displaying data, specifically, in the functional ROIs and motor-related ROIs: mean accuracies were for disgust 57-62%, for anger 62-65%, and for joy 62-69% (Fig 3a).

## Discussion

Our main finding was that expression-specific neural codes are shared between displaying and observing specific emotional facial expressions. Classifier trained on haemodynamic data acquired while subjects displayed different facial expressions successfully predicted neural activation patterns triggered while viewing each of those expressions displayed by unfamiliar models. Highest cross-modal classification accuracies were observed in functional ROIs derived from the experimental data, and in primary motor and somatosensory cortices. Altogether these results support the embodied emotion recognition model, and show that automatically activated, expression-specific neural signatures in sensorimotor and face perception regions of the brain, as well as in emotion circuit ROIs, could support facial expression recognition.

### Discrete neural basis of viewing and displaying facial expressions

We first established that displaying different facial expressions was associated with discrete neural activation patterns, similarly as has been previously demonstrated for hand actions (Dinstein et al. 2008; Smirnov et al 2017). As expected, highest regional classification accuracies were found in the motor and somatosensory cortices. However, accurate classification of the displayed facial expressions was also possible in the regions involved in visual (inferior occipital and fusiform cortices; STS) and affective (amygdala, insula, thalamus, ACC) analysis of facial expression (Haxby et al. 2000). These data thus suggest that displaying emotional facial expressions involved engagement of their affective and visual representations in the brain, likely resulting in visual-motor and affective-motor integration during volitional generation of facial expressions.

In line with previous work (e.g. Said et al. 2010; Peelen et al. 2010; Harry et al. 2013; Wegrzyn et al. 2015), we also confirmed that viewing the facial expressions was associated with distinct expression-specific activation patterns, particularly in the fusiform and inferior occipital cortices, V5, and STS. However, seen expressions could also be successfully decoded from regional activation patterns in the somatosensory (see also Kragel & LaBar, 2016) and motor cortices (see also Liang et al., 2017), and components of the emotion circuit (amygdala, ACC), suggesting that expression-specific affective and somatomotor codes are also activated during facial expression perception.

In general, observing versus displaying facial expressions triggered more widespread brain activation mainly extending to inferior occipital cortex and medial temporal lobe. Despite this, within-modality classification was, in general (with the exception of inferior occipital cortex, V5 and ACC), significantly more accurate for displaying versus observing emotions in all ROIs. Moreover, observed expressions could not be successfully classified from insula and thalamus, suggesting stronger limbic involvement while displaying versus seeing expressions. Altogether these data suggest that the expression-specific neural codes are significantly more discrete when actually generating the expressions (and possibly experiencing the emotion), than when the observers decode the seen facial expressions. Importantly, none of the ROIs alone surpassed accuracy of the whole-brain classification of seen or displayed facial expressions. This suggests that distributed cerebral activation patterns contain the most accurate neural representation of the seen / displayed facial expression. In general sense, these data support the notion that different emotions have discrete neural bases (Kragel & LaBar, 2015; Saarimäki et al. 2016; Wager et al., 2015; Panksepp 1982; Ekman 1992) also in visual and somatomotor domains.

### Shared somatomotor signatures for displayed and observed facial expressions

Both displaying and viewing facial expressions triggered overlapping activity in motor (motor strip and SMA) and somatosensory (S2) cortices. Activation patterns within these regions could be used to predict which facial expression the participants had displayed or viewed, and critically, classifier trained with activation patterns elicited by displaying facial expressions could successfully predict which facial expressions the participants saw in the expression observation condition. Such high cross-modal classification accuracy in primary motor and somatosensory cortices suggests strong embodied component in facial expression recognition.

These data agree with previous neuroimaging studies, which have found common activation patterns in motor and somatosensory cortices during observing and producing facial expressions (Carr et al. 2003; Hennenlotter et al. 2005; Kircher et al. 2013; van der Gaag et al. 2007). These regions also synchronize across a group of individuals seeing or hearing emotional episodes (Nummenmaa et al. 2012; 2014), possibly providing means for shared somatomotor representations of emotions. Furthermore, damage to somatosensory cortex (Adolphs et al. 2000) and its deactivation with TMS (Pourtois et al. 2004) impairs facial expression recognition, while insular cortex supports interoceptive awareness (Critchley et al. 2004). Altogether these data support the models of embodied emotion recognition, which propose that perceiving emotion involves perceptual, somatovisceral, and motoric re-experiencing of the relevant emotion in oneself, and one possible mechanism for that is reactivating modality-specific brain areas without actual behavioural output (Niedenthal 2007). Indeed, behavioural studies have provided that a wide variety of emotions are represented in embodied somatosensory format (Nummenmaa et al, 2014; 2017; Volynets et al., in press), and the present study shows how such embodied signatures of emotions can also contribute to recognizing others’ expressions.

In addition to the somatosensory and motor regions, both seen and displayed facial expressions could be successfully decoded from key regions of the emotion circuit including ACC and amygdala. Critically, cross-modal classification (after functional realignment) was also accurate in these regions. Previously it has been established that these components of the emotion circuit contain discrete neural signatures of experienced emotions (Saarimäki et al. 2016; 2018). Accordingly, these nodes of the emotion system (particularly ACC and amygdala) are involved in both displaying and viewing facial expressions, likely due to their central role in generating the specific emotional states on the basis of internal and external signals. Taken together these data support the position that facial expression recognition is, in addition to visual and somatomotor mechanisms, also supported by affective analysis of the facial signals (Calvo & Nummenmaa 2016).

### Neural signatures for displayed and observed facial expressions are anatomically aligned

For cross-modal classification, we employed pattern recognition after functional realignment of the data from the observing and making facial expressions condition. This was used to test whether the expression-specific neural codes during displaying and viewing facial expressions would be similar yet misaligned across modalities. However, we found that cross-modal classification was already almost as accurate before the functional realignment for most of the ROIS. Realignment only led to a modest increase in classification accuracy in select brain regions (inferior occipital cortex, fusiform cortex, whole grey matter, functional ROIs derived from Observing and Displaying data, their union and intersection). This suggests a direct linkage between the neural systems engaged while displaying and observing facial expressions.

In our experiment, brain regions benefitting most from functional realignment were occipito-ventral components of the core system for facial expression recognition – inferior occipital and fusiform cortices (Haxby et al. 2000, 2010) – whereas little improvement was observed in the emotion and somatomotor systems. Within-modalities classification was highly successful in these regions too. We propose that in these regions neural signatures of displaying and observing certain facial expressions are more anatomically misaligned than in motor and somatosensory cortices, but still distinct and expression-specific.

### Limitations

Because the experiment involved a long and complicated multi-session setup (with two long fMRI sessions, separate behavioural testing and a practice session one day before the scans) we only scanned 12 individuals. However, the results were consistent across the tested subjects, with above chance level cross-modal classification in all subjects. Also, we only included three out of the six canonical basic facial expressions (Ekman 1992) into the study; thus, it is not certain if the results generalize to other facial expressions such as those proposed by Cowen and Keltner (in press). Yet, pattern classification work on facial expression recognition (e.g. Said et al 2010) and “hyperclassification” of seen and executed hand actions (Smirnov et al., 2017) suggests that our results should be generalizable.

## Conclusions

We conclude that displaying and observing emotional facial expressions are supported by shared expression-specific neural codes. A shared set of regions activated during observing and displaying emotional facial expressions includes somatosensory and motor cortices, parts of orbitofrontal cortex, temporal pole and parietal operculum, as well as bilateral caudate, right insula, and right thalamus. Classification analysis successfully distinguished between all tested emotions both within and across modalities. Accurate classification in primary motor and somatosensory cortices and STS suggests strong embodied component in facial expression recognition, while limbic regions are also strongly involved in both displaying and observing emotional facial expressions. Taken together, our results support the embodied emotion recognition model, and suggest that expression-specific neural signatures could underlie facial expression recognition.

## Acknowledgements

This work was supported by Centre for International Mobility (grant TM-14-9213 to S.V.); Academy of Finland (grants #294897 and #265917 to L.N.), and European Research Council Starting Grant (#313000 to L.N.). We also thank Marita Kattelus and Tuomas Tolvanen for her help with the data acquisition.

